# The cluster transfer function of AtNEET supports the ferredoxin-thioredoxin network of plant cells

**DOI:** 10.1101/2022.05.12.491709

**Authors:** Sara I. Zandalinas, Luhua Song, Rachel Nechushtai, David G Mendoza-Cozatl, Ron Mittler

## Abstract

NEET proteins are conserved 2Fe-2S proteins that regulate the levels of iron and reactive oxygen species in plant and mammalian cells. Previous studies of seedlings with constitutive expression of AtNEET, or its dominant-negative variant H89C (impaired in 2Fe-2S cluster transfer), revealed that disrupting AtNEET function causes oxidative stress, chloroplast iron overload, activation of iron-deficiency responses, and cell death. Because disrupting AtNEET function is deleterious to plants, we developed an inducible expression system to study AtNEET function in mature plants using a time-course proteomics approach. Here, we report that suppression of AtNEET cluster transfer function results in drastic changes in the expression of different members of the ferredoxin (Fd), Fd-thioredoxin (TRX) reductase (FTR), and TRX network of Arabidopsis, as well as in cytosolic cluster assembly proteins. In addition, the expression of Yellow Stripe-Like 6 (YSL6), involved in iron export from chloroplasts was elevated. Taken together, our findings reveal new roles for AtNEET in supporting the Fd-TFR-TRX network of plants, iron mobilization from the chloroplast, and cytosolic 2Fe-2S cluster assembly. In addition, we show that AtNEET function is linked to the expression of glutathione peroxidases (GPXs) which play a key role in the regulation of ferroptosis and redox balance in different organisms.

**Highlight:** Using proteomics analysis and an inducible expression system, the iron-sulfur cluster transfer function of AtNEET was found to support the ferredoxin-thioredoxin network of Arabidopsis.

## INTRODUCTION

NEET or CISD (<u>C</u>DGSH <u>I</u>ron-<u>S</u>ulfur <u>D</u>omain) proteins are conserved proteins found in mammalian, plants, fungi, and bacteria (Nechushtai *et al*., 2012, 2020; Inupakutika *et al*., 2017; Sengupta *et al*., 2018). They contain the CDGSH (C-X-C-X2-(S/T)-X3-P-X-<u>C-D-G-(S/A/T)-H</u>) 2Fe-2S cluster binding domain and can participate in different cluster and/or electron transfer reactions (Sengupta *et al*., 2018; Mittler *et al*., 2019; Nechushtai *et al*., 2020). While human cells contain three different NEET proteins (mitoNEET, NAF-1, and MiNT, encoded by CISD1-3, respectively), plants contain only one member of the NEET family, known in Arabidopsis as AtNEET (encoded by AT5G51720; Nechushtai *et al*., 2012). AtNEET structure mostly resembles that of mammalian NAF-1 and mitoNEET, and all three proteins function as homodimers anchored to a membrane. In the case of NAF-1 this membrane is the outer endoplasmic reticulum (ER), mitochondria, or the mitochondrial-associated membranes that connect these two organelles, while in the case of mitoNEET and AtNEET it is primarily the outer mitochondria and chloroplast, respectively (Nechushtai *et al*., 2020). Among the most conserved functions of NEET proteins in different organisms is the regulation of iron and reactive oxygen species (ROS) homeostasis in mitochondria of mammalian cells (Sohn *et al*., 2013), or in chloroplasts of plants (Zandalinas *et al*., 2020b). Suppression of NAF-1 or AtNEET protein levels was found to result in an enhanced accumulation of iron and ROS in the mitochondria or chloroplasts respectively, and this effect was linked to the ability of NAF-1 or AtNEET to bind and release their 2Fe-2S clusters (Darash-Yahana *et al*., 2016; Mittler *et al*., 2019; Zandalinas *et al*., 2020b). Of particular importance to our understanding of NEET function in different biological systems are two studies in which a mutated copy of NAF-1 or AtNEET with a high 2Fe-2S cluster stability (H114C of NAF-1, or H89C of AtNEET) was constitutively expressed in wild type cells to block NEET protein cluster transfer function (Darash-Yahana *et al*., 2016; Zandalinas *et al*., 2020b). By forming heterodimers with the native NEET protein, or complete mutant dimers, the mutated NEET copies functioned as dominant-negative inhibitors of NEET protein function, blocking their different cluster transfer reactions (Darash-Yahana *et al*., 2016; Zandalinas *et al*., 2020b). As indicated above, this inhibition resulted in enhanced iron and ROS accumulation in the mitochondria or chloroplast, that subsequently caused plant and animal cell death (Darash-Yahana *et al*., 2016; Zandalinas *et al*., 2020b). Paradoxically, the constitutive expression of H89C in Arabidopsis was associated with the activation of iron deficiency responses in leaves of plants that accumulated high levels of iron (Zandalinas *et al*., 2020b). This finding suggests that AtNEET, and potentially the levels of 2Fe-2S clusters in plants, could play a key role in the iron sensing mechanism of plants (in leaves). In both mammalian and plant cells, suppression of NEET protein levels or stabilization of the 2Fe-2S clusters of NEET proteins resulted, therefore, in the accumulation of iron and ROS in chloroplasts or mitochondria, activation of the oxidative stress response, activation of mechanisms that prevented iron accumulation in organelles, and cell death (Sohn *et al*., 2013; Darash-Yahana *et al*., 2016; Zandalinas *et al*., 2020b).

Because the constitutive suppression of NEET protein function has a deleterious effect on plant or animal cells, we recently used the Dexamethasone (DEX)-inducible system to drive the expression of NAF-1 or its H114C dominant-negative mutant in cancer cells (Karmi *et al*., 2021). This analysis revealed that in addition to enhanced mitochondrial iron and ROS levels, suppression of NAF-1 function in cancer cells resulted in the enhanced expression of thioredoxin interacting protein (TXNIP) which binds thioredoxin (TRX) and induces oxidative stress (Karmi *et al*., 2021). Despite repeated attempts, we could not however find a homolog of TXNIP in the genome of Arabidopsis, leaving this aspect of NEET function in plant cells unknown. To further explore the function of AtNEET in plants, we used in this study the same DEX-inducible expression system (Aoyama and Chua, 1997) to drive the expression of AtNEET, or its mutated dominant-negative copy H89C, in mature transgenic plants. Using this system we conducted a time-course proteomics analysis to track the cellular changes occurring in plant cells following the inducible expression of AtNEET or H89C. Our findings revealed that suppression of AtNEET function resulted in drastic changes in the expression of different members of the ferredoxin (Fd), Fd:TRX reductase (FTR), and TRX network of Arabidopsis, as well as in the expression level of different members of the cytosolic cluster assembly pathway of plants. In addition, the levels of Yellow Stripe-Like 6 (YSL6), a protein involved in the export of iron from the chloroplast or vacuole was elevated, as well as the expression of different proteins involved in chlorophyll degradation and ROS scavenging. Taken together, our findings reveal new roles for AtNEET in regulating the Fd-TFR-TRX network of cells, iron mobilization from the chloroplast, and cytosolic 2Fe-2S cluster assembly. In addition, we show that the function of AtNEET is affecting the expression of several different ROS scavenging proteins including glutathione peroxidases (GPXs) that play a key role in the regulation of ferroptosis and other stress response pathways in different organisms (Distéfano *et al*., 2021; Karmi *et al*., 2021).

## MATERIALS AND METHODS

### Vector construction and generation of transgenic plants

AtNEET (At5G51720) and H89C (Nechushtai et al., 2012; Zandalinas *et al*., 2020b) cDNAs were amplified by PCR and cloned into the glucocorticoid-inducible transformation pTA7002 vector using XhoI and SpeI sites (Aoyama and Chua, 1997; Supplementary Fig. S1). *Agrobacterium tumefaciens* strain GV3101 was transformed with both constructs and used to obtain DEX-induced AtNEET- and H89C-overexpressing lines using the floral dip procedure (Zhang *et al*., 2006). At least 10 independent lines were selected using hygromycin resistance and expression of AtNEET or H89C upon DEX treatment was determined by quantitative real-time polymerase chain reaction (RT-qPCR; Fig. 1) as described below. Three independent homozygous lines (T4) from both transgenic lines were selected based on both DEX-induced phenotype and NEET or H89C expression (Figs. 1, 2).

**Fig. 1.**
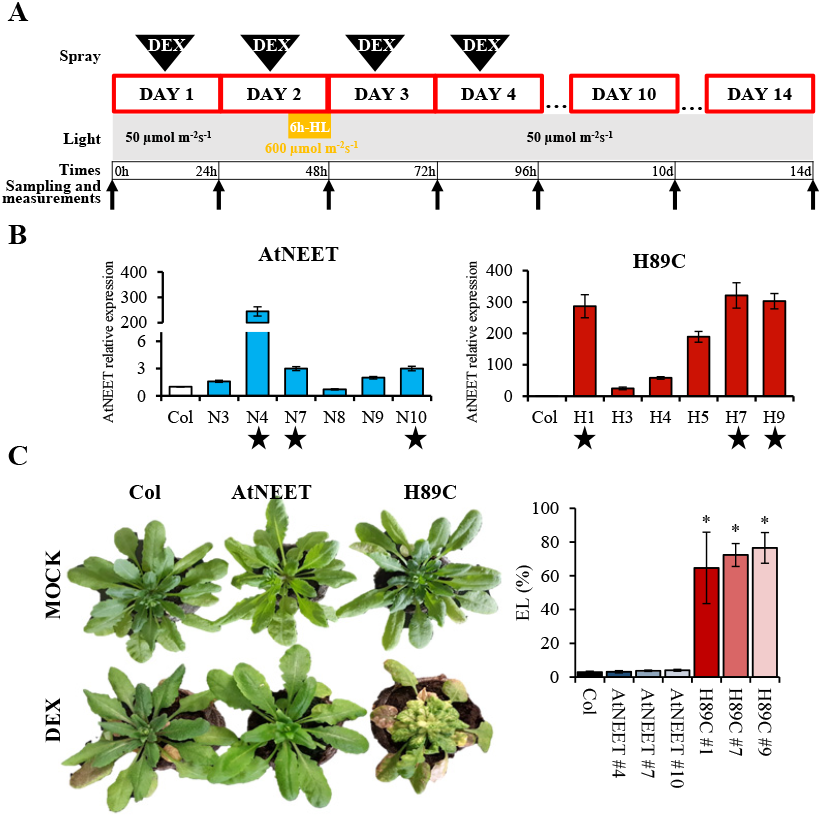
The experimental system used to study the function of AtNEET in Arabidopsis. (A) Outline of the time-course design. Triangular arrows at the top indicate the application of DEX to plants (Col, AtNEET and H89C), and black arrows on bottom indicate the sampling times of all plants for analysis. Yellow box indicates the light stress treatment that was applied on day 2. Please see text for more information. (B) Steady-state transcript expression levels of AtNEET in Col and homozygous AtNEET and H89C plants following 4 doses of DEX application. Stars indicate the plants chosen for further analysis. (C) Representative images of mock and DEX treated Col, AtNEET, and H89C plants on day 14 are shown alongside ion leakage from leaves of the selected lines, also measured on day 14. All experiments were repeated at least three times with similar results. Asterisks denote statistical significance with respect to control (Col) at P < 0.05 (Student t-test, SD, N=5). Abbreviations used: DEX, dexamethasone; EL, electrolyte leakage; HL, high light.

**Fig. 2.**
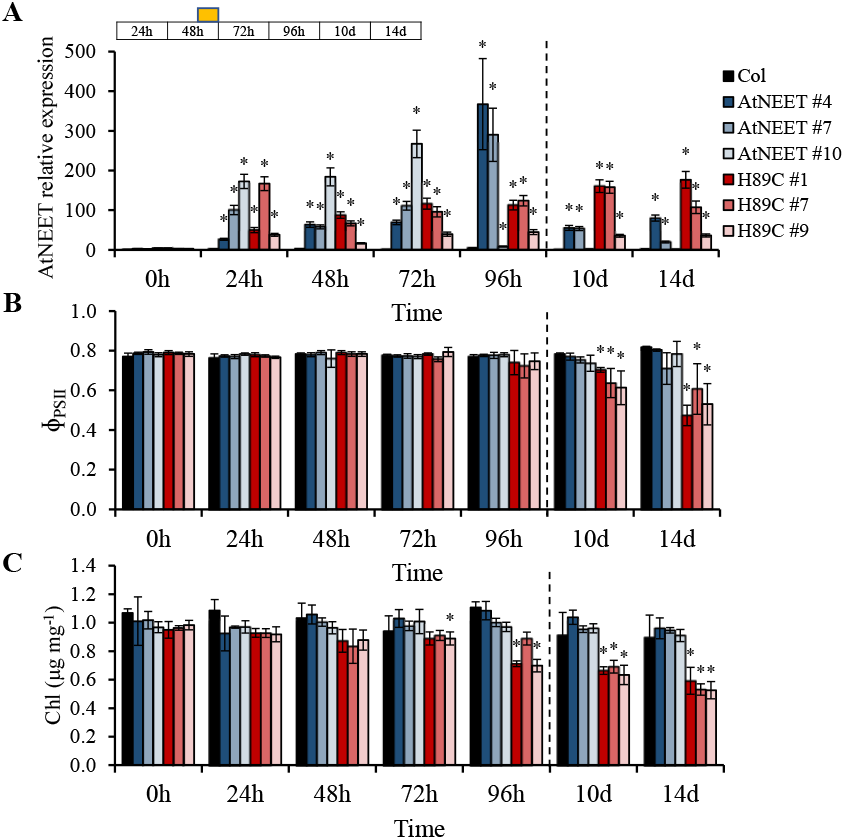
Physiological characterization of Col, AtNEET and H89C plants at the different time points of the experiment. (A) Steady-state transcript expression levels of AtNEET in Col, AtNEET, and H89C plants at the different time points. (B) and (C) Quantum yield of PSII (B) and chlorophyll content (C) measured at the different time points for the different lines. All experiments were repeated at least three times with similar results. Asterisks denote statistical significance with respect to control (0 h) at P < 0.05 (Student t-test, SD, N=5). Abbreviations used: Chl, chlorophyll; PSII, photosystem II.

### Growth conditions and DEX treatment

Col plants and inducible AtNEET and H89C lines were grown in peat pellets (Jiffy-7, Jiffy; http://jiffygroup.com/en/) at 23 °C under long day growth conditions (16-h light/8-h dark, 50 μmol m^-2^ s^-1^). To induce AtNEET or H89C expression, 15-day-old plants were sprayed with a 30 μM DEX (Sigma) solution containing 0.01% (w/v) Tween 20 at the same time of day (10 AM) for 4 days (Fig. 1A). After the second DEX treatment at day 2, plants were subjected to a 6 h-high light treatment (600 μmol m^-2^ s^-1^ from 12 to 6 PM). Leaves of each line were collected at time 0 h (before DEX treatment) and at 24 h, 48 h, 72 h, 96 h, 10 d and 14 d at the same time of the day (9 AM; Fig. 1A). Each experiment was repeated at least three times.

### Proteomics analysis

Leaves from at least 5 plants of Col and inducible AtNEET and H89C lines were collected at each time point as described above (Fig. 1A) and ground to a fine powder in liquid nitrogen with a mortar and pestle. Sample processing, mass spectrometry (MS) analysis and protein identification were performed according to (Dahal *et al*., 2016; Karmi *et al*., 2021). Briefly, grounded leaf tissue was thawed directly into a 1:1 mix of phenol and buffer (Tris-saturated phenol, 0.1 M Tris-HCl pH 8.8, 10 mM EDTA, 200 mM DTT, 0.9 M sucrose). Samples were resuspended in urea buffer (6 M urea, 2 M thiourea, 100 mM ammonium bicarbonate, pH 8.0) and protein quantified using EZQ. An equal amount of protein (50 μg) from each sample was digested with trypsin and peptides were cleaned up using C18 100 μL tips (Pierce), lyophilized, and resuspended in 25 μL of 5% acetonitrile (ACN), 0.1% formic acid (FA). Peptides were analyzed by MS as follows: a 1 μL injection was made onto a C8 trap column (ThermoFisher, μ-precolumn – 300 μm i.d. x 5 mm, C8 Pepmap 100, 5 μm, 100 Å) and separated using a 20 cm long x 75 μm inner diameter pulled-needle analytical column packed with Waters BEH-C18, 1.7 μm reversed phase resin. Peptides were separated and eluted from the analytical column with a gradient of ACN at 300 nL min^-1^. The Bruker nanoElute system was attached to a Bruker tims TOF-PRO mass spectrometer via a Bruker Captive Spray source. Liquid chromatography gradient conditions were as follows: initial conditions were 3% B (A: 0.1% FA in water, B: 99.9% ACN, 0.1% FA), followed by 20 min ramp to 17% B, 17-25% B over 33 min, 25-37% B over 16 min, 37-80% B over 7 min, hold at 80% B for 9 min, ramp back (1 min) and hold (6 min) at initial conditions. Total run time was 92 min. MS data were collected in positive-ion data-dependent PASEF mode over an m/z range of 100 to 1700. One MS and ten PASEF frames were acquired per cycle of 1.16 sec. Target MS intensity for MS was set at 10000 counts s^-1^ with a minimum threshold of 2000 counts s^-1^. An ion-mobility-based rolling collision energy was used: 20 to 59 eV. An active exclusion/reconsider precursor method with release after 0.4 min was used. If the precursor (within mass width error of 0.015 m/z) was higher than 4 times the signal intensity in subsequent scans, a second MSMS spectrum was collected. Isolation width was set to 2 m/z (<700m/z) or 3 (800-1500 m/z). For protein identification, the data were searched against TAIR11 using the following parameters: trypsin as enzyme, 2 missed cleavages allowed; 20 ppm mass error on precursor, 0.1 Da mass error on CID MSMS fragments; carbamidomethyl-Cys fixed modification; oxidized-Met, deamidated-N/Q as variable modifications. Data was then filtered as follows: all identified peptides were filtered for p<0.01 false discovery rate. Data was analyzed using a custom R program using a spectrum count threshold of ≥2 in at least three replicates per group (Supplementary Table S1).

### Electrolyte leakage

Leaves of Col plants and inducible AtNEET and H89C lines from time point 14 d (Fig. 1A) were sampled for electrolyte leakage measurements as described in (Zandalinas *et al*., 2020a) with few modifications. Leaves were immersed in 10 mL of distilled water in 50-mL falcon tubes. Samples were shaken at room temperature for 1 h and the conductivity of the water was measured using a conductivity meter. Leaves were then heated to 95 °C using a water bath for 20 min, shaken at room temperature for 1 h and the conductivity of the water was measured again. The electrolyte leakage was calculated as the percentage of the conductivity before heating over that of after heating.

### RT-qPCR analysis

Relative expression analysis by RT-qPCR was performed according to (Zandalinas *et al*., 2016) by using the CFX Connect Real-Time PCR Detection System (Bio-Rad) and gene-specific primers (Supplementary Table S2).

### Photosynthetic parameters

Quantum yield of Photosystem II (Φ_PSII_) of Col and inducible AtNEET and H89C lines was measured using a portable fluorometer (FluorPen FP 110/S, Photon Systems Instruments, Czech Republic) at each time point described above (Fig. 1A). Photosynthetic measurements were taken for at least 5 plants using two leaves per plant for each time point, line, and experimental repeat.

### Chlorophyll measurements

Chlorophyll extraction was performed as described in (Zandalinas *et al*., 2020b). Briefly, about 50–70 mg of leaves from each line were incubated in 5 mL of N,N-dimethylformamide (DMF) at 4 °C in the dark for 7 d. The absorbance of 1 mL of the DMF extraction was read in a spectrophotometer at 603, 647 and 664 nm, using 1 mL of clean DMF as blank.

### Statistical analysis

Statistical analyses were performed by two-tailed Student’s t-test. Results are presented as the Mean ± SD (asterisks denote statistical significance at P < 0.05 with respect to time 0 h or Col plants). One-way ANOVA along with likelihood ratio (LRT) tests (between time points of each genotype) were used to determine statistically significant changes in protein abundance.

## RESULTS

### Inducible expression of H89C in Arabidopsis

To control the expression of AtNEET and H89C, we generated transgenic plants in which the expression of AtNEET or H89C was driven by a DEX-inducible promoter (Supplementary Fig. S1). To study changes in protein levels in control (Col), AtNEET, and H89C plants following DEX-induced AtNEET or H89C expression, we grew plants under controlled growth conditions for 15 days and then treated them with DEX (30 μM) once a day for 4 days (Fig. 1A). To study the impact of a stress treatment on AtNEET function, on day 2 all plants were subjected to a 6-hour light stress treatment of 600 μmol m^−2^ s^−1^. Plants were kept for a total of 14 days from the beginning of the experiment and sampled at times 0, 24, 48 (immediately after the light stress treatment), 72, and 96 hours, and 10 and 14 days (Fig. 1A). As shown in Fig. 1B, AtNEET transcript expression could be induced by DEX to various levels in different homozygous AtNEET or H89C lines on day 4 (Fig. 1A) of the experiment. Based on this analysis, we chose three AtNEET and three H89C lines for further studies (AtNEET lines 4, 7, and 10, and H89C lines 1, 7, and 9; indicated by stars in Fig. 1B). As shown in Fig. 1C, DEX-treated H89C plants displayed abnormal growth, chlorotic appearance, and ion leakage, indicative of injury or cell death, at 14 days, while control and AtNEET plants (DEX- or mock-treated), or mock-treated H89C plants, did not. These findings reveal that the inducible expression of H89C had a deleterious effect on plants (similar to constitutive expression of H89C; Zandalinas *et al*., 2020b), and demonstrated that the experimental system developed could be used to study the function of NEET proteins in Arabidopsis.

To further characterize plants with inducible expression of AtNEET or H89C, we conducted RT-qPCR analysis of AtNEET expression in the selected AtNEET and H89C lines (Fig. 1B) subjected to the treatments shown in Fig. 1A. As shown in Fig. 2A, DEX-induced AtNEET/H89C expression could be detected in the different lines as early as 24 h post the initial application of DEX. Interestingly, enhanced expression of AtNEET/H89C could also be detected in at least 2 out of 3 AtNEET or H89C lines even at 10 and 14 days (Fig. 2A). To test the effect of AtNEET or H89C expression on photosynthetic activity of plants, we measured the quantum yield of PSII (Φ_PSII_) of all lines included in the experiment. As shown in Fig. 2B, a significant decrease in Φ_PSII_ could only be detected in the three H89C lines at days 10 and 14. In contrast, DEX-treated control or AtNEET plants did not display any significant change in Φ_PSII_. To determine the impact of AtNEET or H89C expression on chlorophyll content, we measured chlorophyll levels of all lines included in the experiment. As shown in Fig. 2C, a significant decrease in chlorophyll content was apparent in all H89C plants on days 10 and 14, as well as at 96 h for some of the H89C lines. In contrast, DEX-treated control or AtNEET plants did not display any significant change in chlorophyll content. Based on the analysis shown in Figs. 1 and 2 we chose the AtNEET line #4 and the H89C line #1 for our in-depth proteomics analysis of AtNEET and H89C induced changes in protein abundance.

### Proteomics analysis of AtNEET and H89C plants following DEX application

Using the experimental design shown in Fig. 1A we conducted an untargeted global proteomics analysis of wild type (WT, Col), AtNEET (AtNEET #4) and H89C (H89C #1) plants. For each time point we used three biological repeats of each line, each with a pool of at least 15 plants. Following identification of the different proteins in each time point, their relative level was compared to time 0 h (within each genotype) and a statistical analysis of fold change in abundance compared to time 0 h was conducted (Supplementary Table S1). Because we treated with DEX and sampled the Col, AtNEET, and H89C lines, side-by-side (Fig. 1A), we could compare the changes that occur in AtNEET plants to those that occur in H89C plants, as well as the changes that occur in Col following DEX application, and the changes associated with light stress in Col, AtNEET and H89C plants. Because the only difference between the AtNEET and the H89C proteins is in one amino acid that causes the cluster to become more stable by more than 10-fold (Nechushtai *et al*., 2012) and induced a dominant-negative effect on AtNEET function (Zandalinas *et al*., 2020b), the inducible expression of H89C could be considered an inhibition of AtNEET cluster transfer function, while the inducible expression of AtNEET could be considered as an enhancement of AtNEET cluster transfer function. In this respect it should be noted that compared to AtNEET, the redox potential of the H89C cluster is shifted by nearly 300 mV and becomes more negative (Nechushtai *et al*., 2012). While the cluster transfer function of AtNEET is suppressed, the electron transfer function of AtNEET could therefore be enhanced in the H89C mutant.

As shown in Fig. 3A, global differences in protein abundance between the different lines were primarily apparent at the 96 h and the 10- and 14-day time points (revealed by a PCA analysis of all results combined). This finding was in agreement with changes in Φ_PSII_ and chlorophyll content that were also apparent at these time points (Fig. 2B, C), suggesting that it took time for the inducible expression of H89C and AtNEET to cause an overall change in protein abundance. While changes in AtNEET protein abundance in control (Col) plants were not significant throughout the experiment, the abundance of the AtNEET and H89C proteins was elevated at all time points (Fig. 3B). Interestingly, while the abundance of H89C was stable throughout the entire experiment (about 4-5-fold higher compared to time 0 h), the abundance of AtNEET was much more variant and higher than that of H89C (Fig. 3B). This finding could suggest that due to the toxicity of the H89C protein, its levels were maintained low in cells. This possibility could also explain why it took time for H89C plants to develop a visible phenotype (Fig. 1C) and cause global changes in protein abundance (Fig. 3A).

**Fig. 3.**
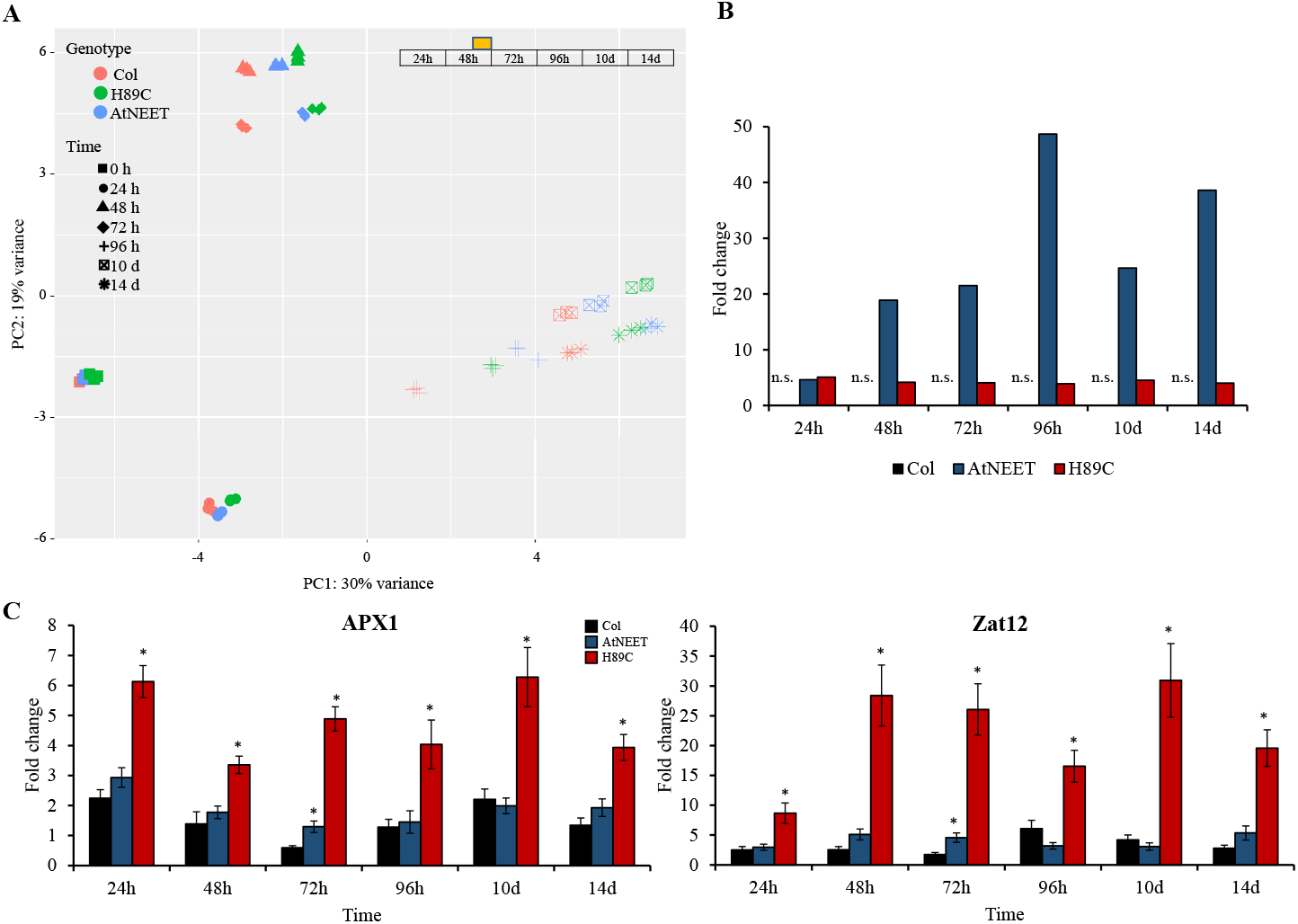
Proteomics analysis and expression measurements of selected transcripts at the different time points. (A) Principal component analysis (PCA) of the proteomics results obtained for the different lines at the different time points of the experiment. (B) Expression level of AtNEET in Col, AtNEET and H89C plants at the different time points. (C) Steady-state transcript expression levels of APX1 and Zat12 in Col, AtNEET, and H89C plants at the different time points. All experiments were repeated at least three times with similar results. Asterisks denote statistical significance with respect to control (Col) at P < 0.05 (Student t-test, SD, N=5). Abbreviations used: APX1, ascorbate peroxidase 1; n.s., not significant; PC, principal component; Zat12, zinc finger protein ZAT12.

Constitutive expression of H89C in seedlings was previously reported to cause oxidative stress and induce the expression of several different ROS-response transcripts (Zandalinas *et al*., 2020b). To determine whether inducible expression of H89C in mature plants would also cause the activation of an oxidative stress response, we used RT-qPCR to measure the transcript expression of two ROS response transcripts, namely *APX1* and *ZAT12* (Davletova *et al*., 2005b,*a*). As shown in Fig. 3C, the expression of *APX1* and *ZAT12* was significantly elevated in H89C plants at all time points. This finding suggests that H89C expression results in enhanced oxidative stress, but that the plant buffering capacity for oxidative stress could shield its metabolism for at least 72 h before changes in chlorophyll, Φ_PSII_, and global protein abundance occur (Figs. 2 B, C, 3A).

To compare the effects of AtNEET or H89C inducible expression in mature plants to those of constitutive AtNEET or H89C expression in seedlings, we compared the large proteomics data sets obtained in this study (Supplementary Table S1) with the proteomics datasets previously obtained for constitutive expression of AtNEET or H89C in seedlings (Zandalinas *et al*., 2020b). As shown in Fig. 4, more that 50% of the proteins previously identified in seedlings with constitutive expression of AtNEET or H89C (Zandalinas *et al*., 2020b) were also identified by our current inducible expression analysis conducted with mature plants. This finding supported the validity of our experimental system and demonstrated that compared to the previous analysis conducted with constitutive expression of AtNEET or H89C, our inducible expression strategy identified many more proteins.

**Fig. 4.**
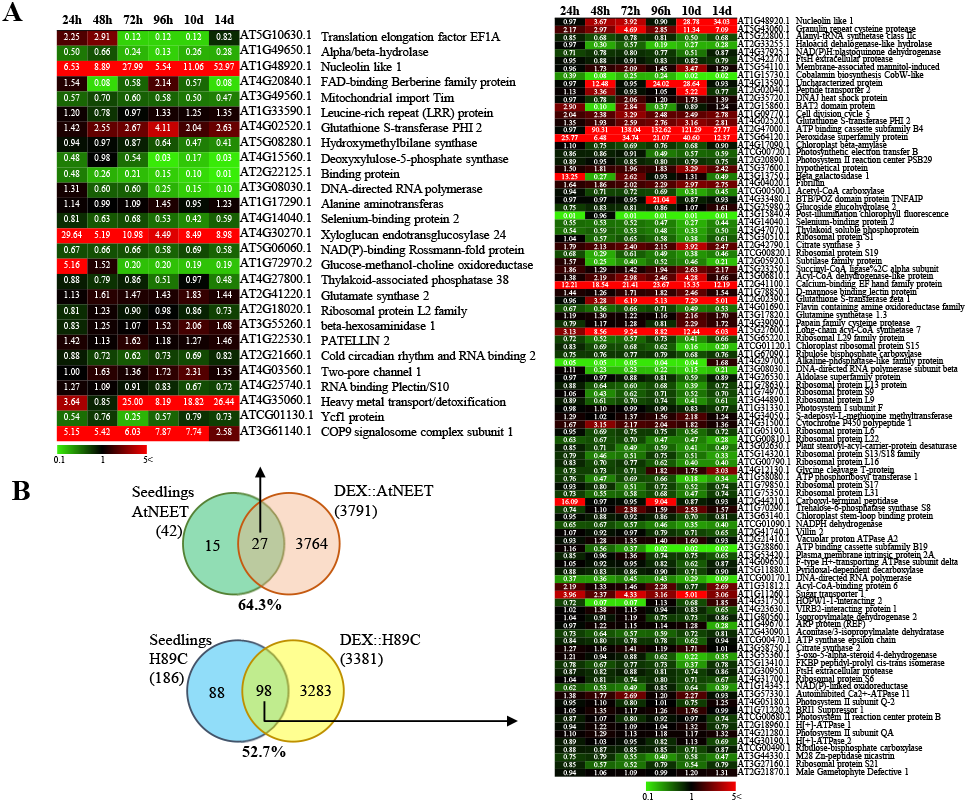
Comparison between the proteomics results obtained with the inducible expression system in this study and the results obtained with constitutive expression of AtNEET and H89C. (A) Heat maps for the expression pattern of proteins shared between the two experimental systems. (B) Venn diagrams showing the overlap between the two experimental systems (inducible expression in mature plants *vs* constitutive expression in seedlings). Proteomics results of constitutive AtNEET and H89C expression were obtained from Zandalinas et al., 2020b. All experiments were repeated at least three times with similar results.

### Altered abundance of different components of the cytosolic iron-sulfur cluster assembly (CIA) pathway in AtNEET and H89C plants following DEX application

We previously reported that the steady state expression of several different transcripts involved in the iron-sulfur biogenesis pathways of the cytosol, chloroplast and mitochondria is altered in plants with constitutive expression level of H89C (Zandalinas *et al*., 2020b). However, whether these changes are directly related to H89C, or an indirect effect of its constitutive expression on plants was unknown. As shown in Fig. 5A, using the inducible expression system, we now report that triggering the expression of AtNEET or H89C results in direct, and in many cases immediate, changes in the abundance of different proteins involved in the CIA pathway. Of particular interest to the function of AtNEET are CIA1, NBP35 and DRE2, especially since AtNEET was found to transfer its clusters to DRE2 (Zandalinas *et al*., 2020b), and DRE2 transfer its clusters to NBP35 that then transfer them to the CIA1-associated complex (Zhang *et al*., 2008; Balk and Pilon, 2011). The changes in transcript expression reported previously in plants with constitutive expression of H89C (Zandalinas *et al*., 2020b) are therefore strengthen and extended now with results from a dynamic inducible system that is coupled to proteomics analysis (Figs. 1A, 5A). Moreover, changes in the abundance of CIA1, that plays a key function in the CIA pathway (Braymer *et al*., 2021), were not previously identified and are therefore a new finding of this study (Fig. 5A). As shown in Fig. 5B, the inducible expression of H89C or AtNEET also resulted in changes in the steady-state transcript levels of CIA1, further supporting its identification by the proteomics analysis. Taken together, the results shown in Fig. 5 support our previous study that used constitutive expression of AtNEET and H89C in seedlings (Zandalinas *et al*., 2020b) and reveal that CIA1 is directly responding to changes in AtNEET function in mature plants.

**Fig. 5.**
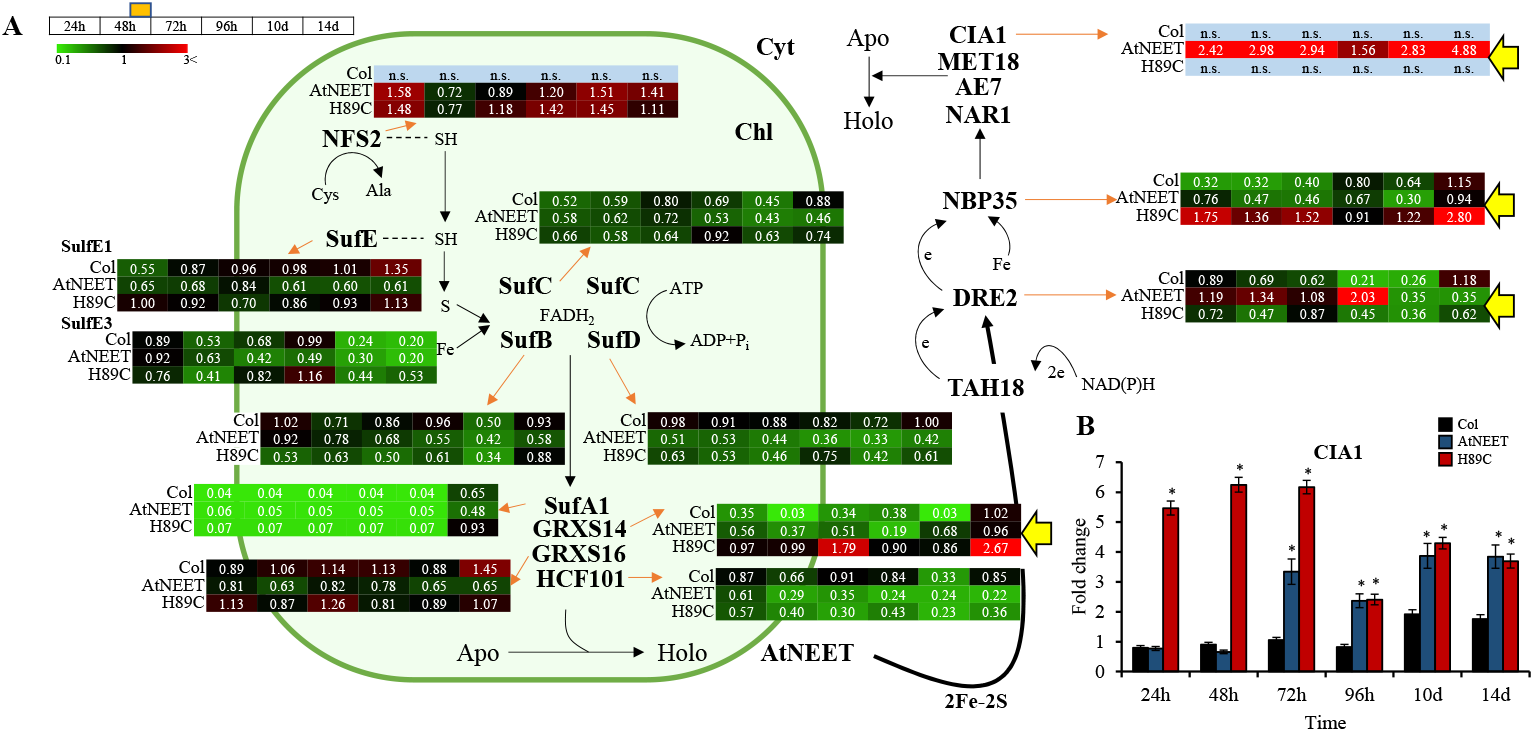
Changes in protein and transcript expression associated with iron-sulfur cluster assembly in the chloroplast and cytosol during the course of the experiment. (A) Pathway and heat maps for the expression pattern of different proteins with a significant change in expression (in at least one time point, compared to time 0 h within each genotype) belonging to the iron-sulfur cluster assembly of Arabidopsis at the different time points. (B) Steady-state transcript expression levels of CIA1 in Col, AtNEET, and H89C plants at the different time points. Yellow arrows highlight proteins of interest. All experiments were repeated at least three times with similar results. Asterisks denote statistical significance with respect to control (Col) at P < 0.05 (Student t-test, SD, N=5). Abbreviations used: AE7, AS1/2 Enhancer 7; Apo, apo-protein; Chl, chloroplast; CIA1, Cytosolic Iron-Sulfur Protein Assembly 1; Cyt, cytosol; DRE2, Homolog of Yeast DRE2; e, electron; GRXS14, Glutaredoxin S14; GRXS16, Glutaredoxin S16; HCF101, High-Chlorophyll-Fluorescence 101; Holo, holo-protein; MET18, Homolog of Yeast MET18; NAR1, Homolog of Yeast NAR1; NBP35, Nucleotide Binding Protein 35; NFS2, Nifs-Like Cysteine Desulfurase 2; n.s., not significant; SufA1, Sulfur A1; SufB, Sulfur B; SufC, Sulfur C; SufD, Sulfur D; SufE, Sulfur E; TAH18, diflavin reductase.

### Altered abundance of iron efflux proteins following alterations in AtNEET function

We previously reported that constitutive expression of H89C resulted in the accumulation of iron in chloroplasts that was paradoxically coupled with transcriptional activation of leaf iron deficiency responses (Zandalinas *et al*., 2020b). However, whether this response was also accompanied by changes in the expression of different iron transport proteins was unknown. As shown in Fig. 6, using the inducible expression system, we found that triggering the expression of H89C in mature plants results in an early and strong induction in the abundance of the chloroplastic (and potentially also vacuolar) iron export protein YSL6 (Divol *et al*., 2013; Conte *et al*., 2013). In contrast, abundance of the chloroplastic iron import protein PIC1 (Duy *et al*., 2007, 2011) primarily increased following inducible expression of AtNEET (Fig. 6). Interestingly, the expression level of transcripts encoding YSL6 or PIC1 did not significantly change in transgenic seedlings with constitutive expression of AtNEET or H89C (Zandalinas *et al*., 2020b). Our new finding that the abundance of the iron export protein YSL6 is rapidly enhanced in H89C leaves upon DEX treatment supports our previous findings that chloroplasts of H89C seedlings with constitutive expression of H89C accumulate high levels of iron (and therefore enhance the abundance of YSL6; Fig. 6), while activating iron deficiency responses (Zandalinas *et al*., 2020b).

**Fig. 6.**
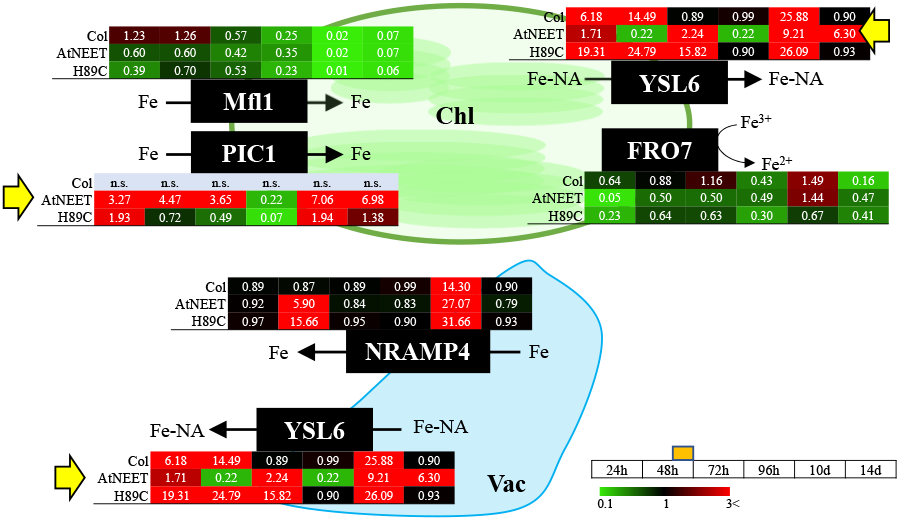
Changes in protein expression levels associated with iron/metal transport during the course of the experiment. Pathway and heat maps for the expression pattern of different proteins with a significant change in expression (in at least one time point, compared to time 0 h within each genotype) associated with the transport of iron and other metals into and out of the chloroplast and vacuole are shown. Yellow arrows highlight proteins of interest. All experiments were repeated at least three times with similar results. Abbreviations used: Chl, chloroplast; FRO7, Ferric Reduction Oxidase 7; Mfl1, Mitoferrin-like 1; NRAMP4, Natural Resistance Associated Macrophage Protein 4; n.s., not significant; PIC1, Permease In Chloroplasts 1; Vac, vacuole; YSL6, Yellow Stripe Like 6.

### Alterations in the Fd-FTR-TRX network of Arabidopsis following the inducible expression of AtNEET or H89C

In a previous study we demonstrated that inducible expression of the NAF-1 H114 mutant (with a high 2Fe-2S cluster stability) in cancer cells results in the enhanced expression of TXNIP that binds TRXs and induces oxidative stress and ferroptosis (Karmi *et al*., 2021). Although plants were not found to have a clear homolog of TXNIP, they contain an extended network of TRXs that is in many instances linked to Fd via FTRs (*e.g*., Kang *et al*., 2019; Balsera and Buchanan, 2019; Cejudo *et al*., 2021; Ojeda *et al*., 2021; Fig. 7A). AtNEET was originally identified as an 2Fe-2S cluster donor to Fd1 (Nechushtai *et al*., 2012), suggesting that alterations in AtNEET cluster transfer function could cause alterations in the entire Fd-FTR-TRX network. As shown in Fig. 7A, the inducible expression of AtNEET or H89C indeed caused drastic changes in the abundance of different Fds, FTRs, and TRXs. Examples for these changes include the TRX AT1G21350 that was specially upregulated, the TRX AT1G76020 that was specifically downregulated, and FdC1 that was downregulated following H89C induction plants, and Fd1 and the 2Fe-2S Fd-like protein AT4G32590 that were upregulated following AtNEET induction in plants. As shown in Fig. 7B, the steady state level of transcripts encoding some of these proteins was also significantly altered following the inducible expression of AtNEET or H89C. The expression of *TPX1* and *TPX2* was also found to be upregulated in our previous transcriptomics data set of plants with constitutive expression of H89C (Supplementary Fig. S2). Taken together, the findings presented in Fig. 7 support a role for AtNEET in modulating and supporting the TRX network of Arabidopsis, possibly through providing 2Fe-2S clusters to Fd.

**Fig. 7.**
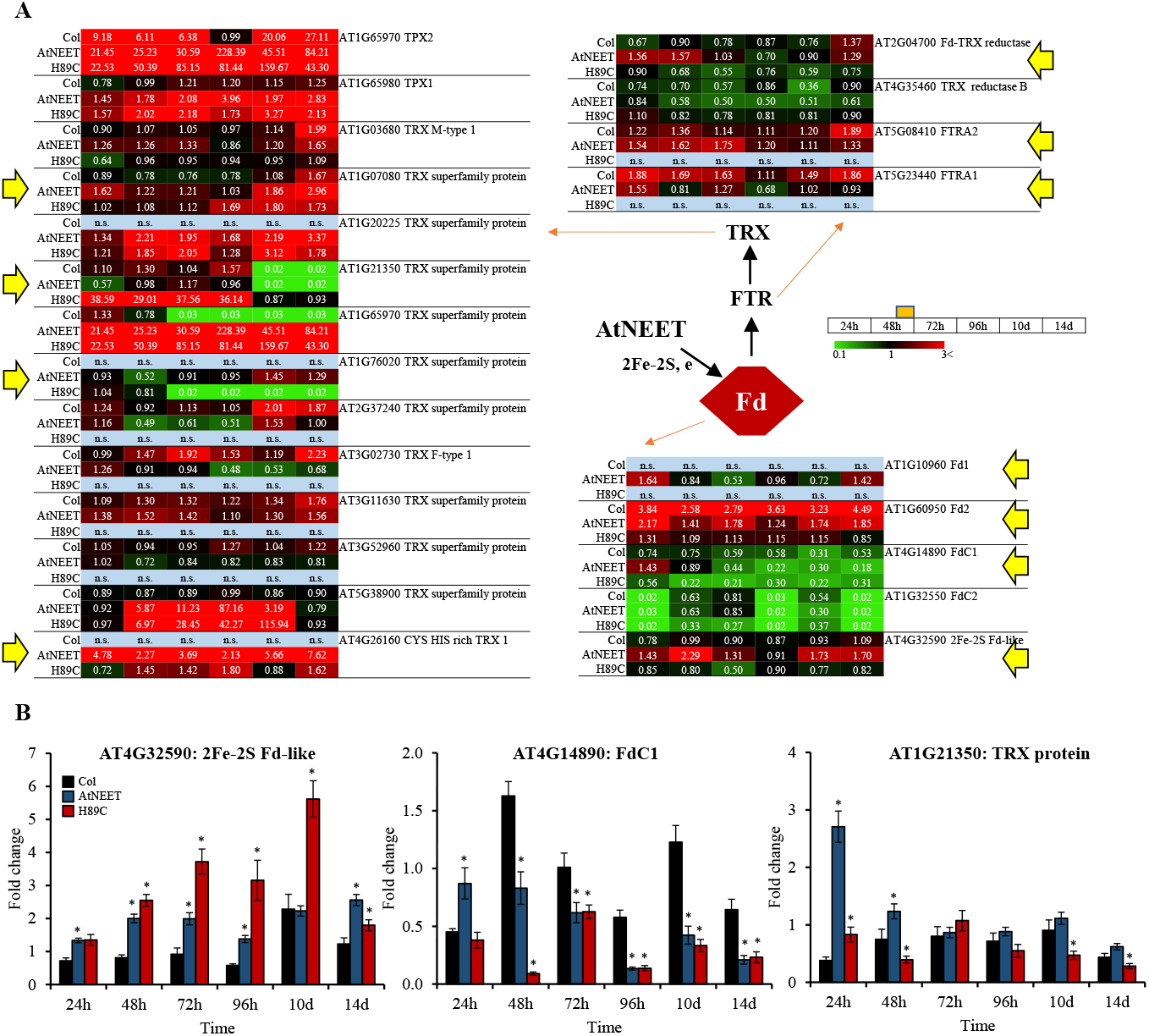
Changes in protein and transcript expression associated with the ferredoxin (Fd), Fd-thioredoxin (TRX) reductase (FTR) and/or TRX during the course of the experiment. (A) Pathway and heat maps for the expression pattern of different proteins with a significant change in expression (in at least one time point, compared to time 0 h within each genotype) associated with the Fd-FTR-TRX network of Arabidopsis at the different time points. (B) Steady-state transcript expression levels of an 2Fe-2S Fd-like, FdC1 and a TRX protein in Col, AtNEET, and H89C plants at the different time points. Yellow arrows highlight proteins of interest. All experiments were repeated at least three times with similar results. Asterisks denote statistical significance with respect to control (Col) at P < 0.05 (Student t-test, SD, N=5). Abbreviations used: Fd, ferredoxin; FTR, ferredoxin-thioredoxin reductase; FTRA1, Ferredoxin/thioredoxin reductase subunit A1; FTRA2, Ferredoxin/thioredoxin reductase subunit A2; n.s., not significant; TRX, thioredoxin; TPX, thioredoxin-dependent peroxidase.

Because Fds transfer electrons to many different essential proteins in plant cells, we also studied the abundance of additional proteins that serve as electron acceptors of Fds. As shown in Supplementary Fig. S3, the abundance of pheophorbide A oxygenase (PAO), a Rieske-type iron– sulfur protein involved in chlorophyll degradation, was rapidly upregulated in H89C plants. Chlorophyll degradation is also part of the initial Fe deficiency response to prevent the accumulation of toxic tetrapyrrole intermediaries. In addition, the abundance of the nitrogen stress related protein glutamine synthetase 2 (GS2) was upregulated in AtNEET plants. These findings extended the list of cellular pathways potentially supported by AtNEET to include chlorophyll catabolism and nitrogen metabolism.

### Changes in the abundance of different ROS scavenging enzymes following alterations in AtNEET function

The TRX network is directly linked to the function of different proteins that scavenge ROS such as H_2_O_2_ (*e.g*., through GPX or the TRX-peroxiredoxin cycles; Balsera and Buchanan, 2019; Foyer *et al*., 2020; Meyer *et al*., 2020). Because the inducible expression of AtNEET had such a dramatic effect on the Fd-FTR-TRX redox network of Arabidopsis (Fig. 7), we tested the abundance of different proteins involved in H_2_O_2_ scavenging (Willems *et al*., 2016). As shown in Fig. 8, the abundance of GPX5 and GPX6, monodehydroascorbate reductase (MDAR), and three ascorbate peroxidases (APX1, APX6 and stromal APX) was altered following the inducible expression of AtNEET or H89C. The expression of transcripts encoding *GPX6* and *stromal APX* was also found to be upregulated in our previous transcriptomics data set of plants with constitutive expression of H89C (Supplementary Fig. S2). The findings described above are particularly interesting since previous studies found that the mammalian NEET protein mitoNEET can be reduced by glutathione (GSH) or GSH reductase (GR), as well as oxidize H_2_O_2_ (Landry and Ding, 2014; Landry *et al*., 2015). In addition to altering the abundance different components of the Fd-FTR-TRX network (Fig. 7), the disruption in AtNEET function could therefore also impair the H_2_O_2_ metabolizing capacity of cells.

**Fig. 8.**
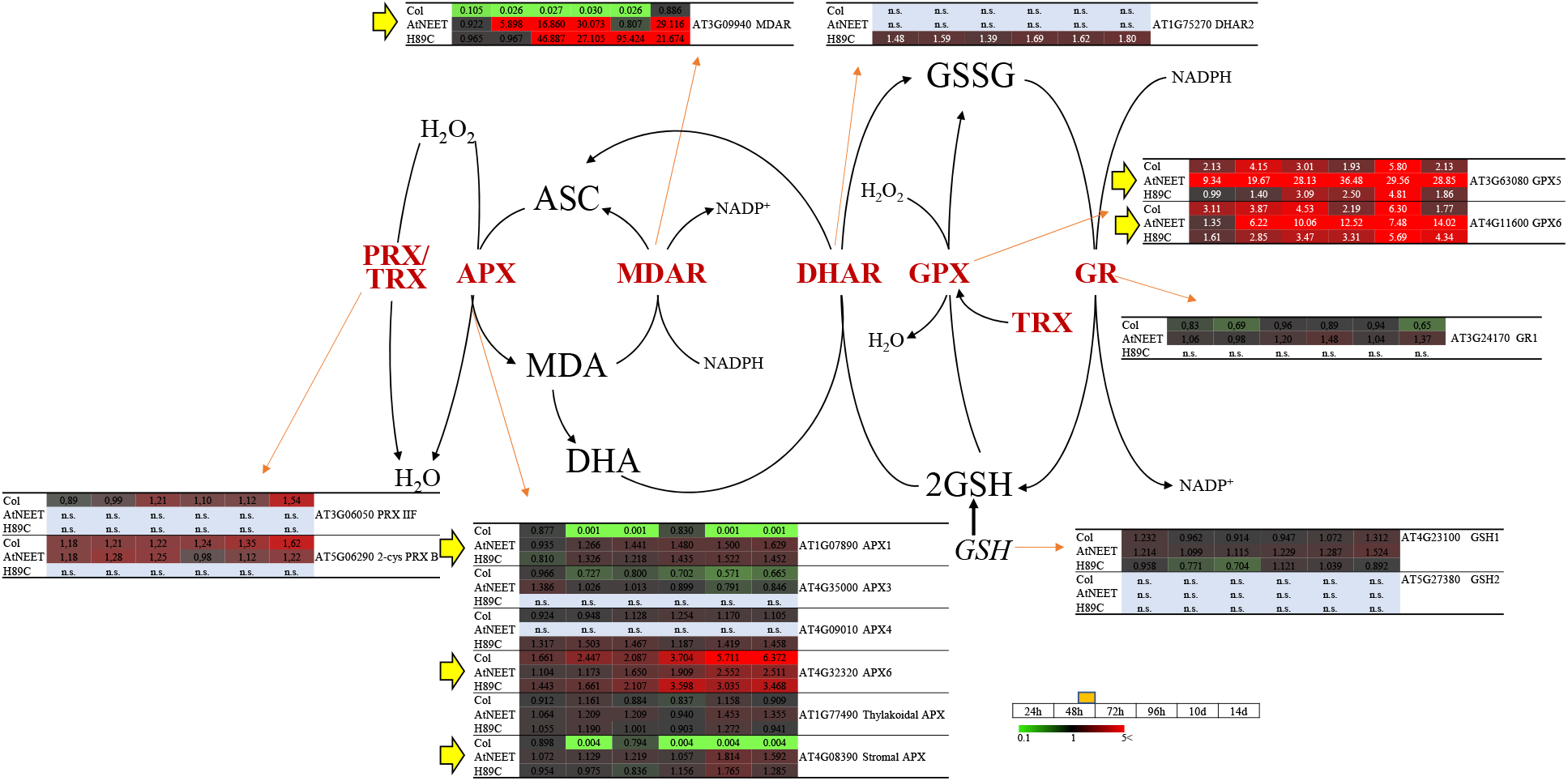
Changes in protein expression associated with reactive oxygen species (ROS) scavenging during the course of the experiment. Pathway and heat maps for the expression of different proteins with a significant change in expression (in at least one time point, compared to time 0 h within each genotype) associated with ROS scavenging in Arabidopsis at the different time points are shown. Yellow arrows highlight proteins of interest. All experiments were repeated at least three times with similar results. Abbreviations used: APX, ascorbate peroxidase; ASC, ascorbate; DHA, dehydroascorbate; DHAR, dehydroascorbate reductase; GPX, glutathione peroxidases; GR, glutathione reductase; GSH, glutathione; GSH1, glutamate-cysteine ligase 1; GSH2, glutathione synthase 2; GSSG, oxidized glutathione; MDA, monodehydroascorbate; MDAR, monodehydroascorbate reductase; n.s., not significant; PRX; peroxiredoxin; TRX, thioredoxin.

Because our time course proteomics analysis was conducted with only one selected H89C line, we tested changes in the expression of selected transcripts in all three H89C lines. As shown in Supplementary Fig. S4, changes in the expression of *Fd, DRE2, SufB, SufD*, and *GRSX14* were similar between all three lines at day 4.

## DISCUSSION

Studying gene function using constitutive gain- or loss-of-function mutants is a powerful approach. However, it has the drawback that the altered gene function exists from the very first stage of the organism (mutant) development. In cases in which altering the gene function has deleterious effects, such as in the case of the H89C mutant of AtNEET (Zandalinas *et al*., 2020b), the study of gene function at a mature stage of the organism might not even be possible. To address this problem and to study AtNEET function in mature plants, we used an inducible expression system. This system allowed us to observe dynamic changes in protein abundance resulting from disrupting the cluster-transfer function of AtNEET in cells. When the function of two pathways or proteins is coupled in cells, altering the function of one of them could cause the elevated or suppressed expression or abundance of the other, depending on the nature of the regulatory circuit that controls the expression of the pathway (*e.g*., negative or positive feedback loops). For this reason, we considered each significant change in protein abundance, observed between different time points in AtNEET and/or H89C plants following DEX application in our experiments (up- or down-regulated), as evidence for a potential link to AtNEET function.

To further address the function of AtNEET under altered environmental conditions, and to place the biological systems linked to AtNEET under strenuous conditions, we subjected all plants studied to a light stress treatment at day two following DEX application (Fig. 1A). As shown in Supplementary Fig. S3, for FNR expression, this treatment affected all plant lines studied (WT, H89C and AtNEET). However, compared to WT, it had a more significant effect on protein abundance in the H89C and/or AtNEET, as shown for example in Fig. 7A for TPX1, as well as many other examples discussed below. Changes that occurred within the first 24 h were therefore related to the DEX induced alterations in AtNEET or H89C expression, while changes that occur at 48 h and onwards were changes that occurred due to the DEX induced alterations in H89C or AtNEET expression, as well as the light stress treatment. Overall, there was a good overlap between changes in protein abundance identified by the current proteomics analysis conducted with inducible expression of AtNEET and H89C, and the previous study that used constitutive expression of these proteins (*e.g*., Fig. 4; Zandalinas *et al*., 2020b). In addition, the inducible expression of H89C had deleterious effects on plant growth, chlorophyll content and cell integrity, as evident by the visible phenotype, ion leakage measurements and chlorophyll content (Figs. 1, 2). However, as discussed below, compared to the constitutive expression of AtNEET or H89C, the dynamics nature of the current experimental design allowed us to identify additional and/or new clues to AtNEET function in plants and revealed potential new links between AtNEET and different metabolic and acclimation networks in plants.

We previously reported that the expression of several transcripts encoding chloroplastic and cytosolic Fe-S cluster assembly proteins is upregulated in plants with constitutive expression of H89C (Zandalinas *et al*., 2020b). In addition, we reported that the expression level of several FeS proteins is suppressed in H89C plants, and that AtNEET can transfer its clusters to DRE2 that is a member of the CIA complex in Arabidopsis (Zandalinas *et al*., 2020b). However, whether the expression of different CIA proteins is altered in response to altering the function of AtNEET was unknown. Here we show for the first time that the protein expression of CIA1 and DRE2 is upregulated upon inducible expression of AtNEET, but not H89C, suggesting that augmenting the level of AtNEET results in higher expression of some CIA proteins (Fig. 5). Taken together with our previous transcriptomics analysis (Zandalinas *et al*., 2020b), our findings, shown in Fig. 5, support a model in which AtNEET plays a central role in transferring clusters from within the chloroplast to the cytosol and that altering the cluster transfer ability of AtNEET impairs this process (Fig. 9). In this respect it should be noted that several recent studies support a similar function for mammalian NEET proteins, forming a cluster transfer relay between the mitochondria and the cytosol. In this new role, MiNT (that is not found in plants) transfers its clusters to mitoNEET through the VOLTAGE-DEPENDENT ANION CHANNEL (VDAC) channel, that then transfers its clusters to NAF-1 and Anamorsin (a component of the mammalian CIA complex and a homolog of the plant DRE2 protein; Lipper *et al*., 2015; Karmi *et al*., 2017, 2022). Although the chloroplast is not known to contain VDAC, and NEET proteins are represented in Arabidopsis by only one gene member (AtNEET), it appears that transferring clusters from within an organelle (mitochondria in mammalian and chloroplast in plants) to the cytosol (to the CIA pathway) is a conserved function of NEET proteins.

**Fig. 9.**
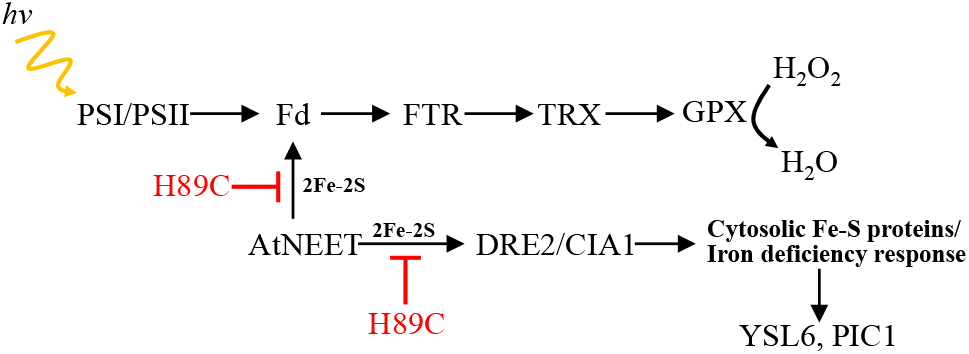
A simplified model for the dual role of AtNEET in plants. By providing 2Fe-2S clusters to ferredoxins, AtNEET is shown to support the function of the ferredoxin (Fd), Fd-thioredoxin (TRX) reductase (FTR), and TRX network of Arabidopsis (top). In addition, AtNEET is shown to play a key role in the mobilization of 2Fe-2S clusters from within the chloroplast to the cytosol and this function is shown to be important for regulating the level of different Fe-S cluster-containing proteins as well as the iron deficiency response of Arabidopsis. Functioning as a dominant-negative inhibitor of AtNEET iron cluster transfer functions, H89C is shown to block these two pathways. The model shown was developed based on the results obtained in the current study and the results presented in Zandalinas et al., 2020b. Abbreviations used: CIA1, Cytosolic Iron-Sulfur Protein Assembly 1; DRE2, Homolog of Yeast DRE2; Fd, ferredoxin; FTR, ferredoxin-thioredoxin reductase; GPX, glutathione peroxidase; PIC1, Permease In Chloroplasts 1; PSI, photosystem I; PSII, photosystem II; TRX, thioredoxin; YSL6, Yellow Stripe Like 6.

We previously proposed that suppressing the cluster transfer activity of AtNEET via constitutive expression of H89C activates the iron deficiency response of Arabidopsis, potentially due to enhanced accumulation of iron in the chloroplast that is coupled with decreased availability of FeS clusters in the cytosol (Zandalinas *et al*., 2020b). This model was proposed based on changes the expression level of several transcripts involved in the iron deficiency response of Arabidopsis, as well as changes in the expression of different transcripts involved in iron efflux from the chloroplast. However, the impact of suppressing AtNEET cluster transfer function on the expression level of different proteins involved in these pathways was unknown. Here we report for the first time that the protein expression of YSL6, involved in the export of iron from the chloroplast (and potentially the vacuole) to the cytosol (Divol *et al*., 2013; Conte *et al*., 2013), is rapidly and strongly enhanced following H89C induction (with some induction at early and late time points following AtNEET induction; Fig. 7). In addition, we report that the protein expression level of PIC1, involved in iron uptake into chloroplasts (Duy *et al*., 2007, 2011), is primarily enhanced upon induction of AtNEET expression (with some induction at early and late time points upon H89C induction; Fig. 7). Taken together with our transcriptomics analysis (Zandalinas *et al*., 2020b), the findings presented in Fig. 7 support the proposed involvement of AtNEET in iron metabolism in plant cells and demonstrate for the first time that changes in AtNEET cluster transfer function translate into changes in the expression of proteins involved in the mobilization of iron from and to the chloroplast (Fig. 9).

We previously demonstrated that AtNEET can transfer its clusters to Fd1 (Nechushtai *et al*., 2012). However, the biological significance of this cluster transfer reaction was unknown. Here we demonstrate for the first time, that upon suppression of AtNEET cluster transfer function, major alterations occur in the protein abundance of different Fds, FTRs and TRXs (Fig. 6). Thus, while the abundance of Fd1, Fd2, FdC1, and an 2Fe-2S Fd was either suppressed or unchanged upon induction of H89C expression, the abundance of Fd1, Fd2, and the 2Fe-2S Fd-like protein was mostly enhanced upon induction of AtNEET expression (Fig. 6). A similar pattern was observed for at least three FTRs (Fd-TRX reductase, FTRA1, and FTRA2). In contrast, TRXs displayed a more variable response with some TRXs upregulated in H89C (*e.g*., AT1G21350) and some suppressed (*e.g*., AT1G76020). Alterations in AtNEET cluster transfer function could therefore be associated with significant changes in the Fd-FTR-TRX network and this finding could be explained by a deficiency in the ability of AtNEET to donate its clusters to Fd (Nechushtai *et al*., 2012; Fig. 9). If AtNEET is prevented from transferring its clusters to Fds (via *e.g*., H89C expression), the entire Fd-FTR-TRX could therefore be affected, resulting in drastic changes in the cells’ redox states and thereby in many cellular functions. In support of this possibility is also the reduced expression of transcripts encoding Fd1 upon DEX induction of H89C (Supplementary Fig. S4). AtNEET could therefore be supporting the Fd-FTR-TRX network by keeping Fd supplied with 2Fe-2S cluster, maintaining its activity.

Because some plant GPXs are thought to utilize TRXs for their reduction/oxidation cycles (Maiorino *et al*., 2015; Meyer *et al*., 2020) that would directly control the levels of H_2_O_2_, as well as the redox regulation of many proteins in plants, the suppression of Fd function upon AtNEET cluster transfer inhibition could also cause the induction of an oxidative stress response (also shown by *Zat12* and *APX1* induction in Fig. 3C). Indeed, the abundance of two GPXs was found to be significantly enhanced upon induction of AtNEET expression (Fig. 8), supporting a link between AtNEET and GPX expression (Fig. 9). In this respect it should be noted that in mammalian cells GPXs are thought to regulate the process of ferroptosis (Jiang *et al*., 2021), and that we recently reported that suppressing the cluster transfer function of NAF-1 (via inducible expression of H114C) altered GPX expression, activated ferroptosis, and caused the enhanced accumulation of TXNIP (a major regulator of the mammalian TRX network) in cancer cells (Karmi *et al*., 2021). Taken together, our findings in plant and mammalian cells reveal a potentially new and conserved role for NEET proteins in regulating the TRX network of cells, as well as suggest that AtNEET could play a role in ferroptosis activation in plant cells (Zandalinas *et al*., 2020b; Distéfano *et al*., 2021; Karmi *et al*., 2021). In the context of this potential new role for AtNEET in supporting the Fe-FTR-TRX network and GPX function by providing clusters to Fd (Fig. 9), it is worth mentioning that previous studies conducted with the mammalian mitoNEET protein revealed that this protein interacts with glutathione reductase (GR), can accept electrons from glutathione and can oxidize H_2_O_2_ (Landry and Ding, 2014; Landry *et al*., 2015). Based on these findings it was proposed that mitoNEET could function as a sensor or scavenger of ROS. While a similar function was not reported for AtNEET, our findings that suppressing the cluster transfer function of AtNEET causes oxidative stress in plants (Zandalinas *et al*., 2020b; Fig. 3C), might support a similar function for AtNEET in plants. The nature of the interactions between AtNEET and the Fd-FTR-TRX and/or the glutathione/GR/GPX networks requires further studies, especially since NEET proteins can transfer or accept clusters, as well as electrons, to or from other cellular proteins (Zuris *et al*., 2011; Nechushtai *et al*., 2012; Landry and Ding, 2014; Landry *et al*., 2015; Li *et al*., 2018; Tasnim *et al*., 2020).

## Abbreviations

CIA: cytosolic iron-sulfur cluster assembly
CISD: CDGSH Iron-Sulfur Domain
DEX: dexamethasone
Fd: ferredoxin
FTR: ferredoxin:thioredoxin reductase
GSH: glutathione;
GPX: glutathione peroxidase
GR: glutathione reductase
MDAR: monodehydroascorbate reductase
MS: mass spectrometry
ROS: reactive oxygen species
TRX: thioredoxin
TXNIP: thioredoxin interacting protein

## SUPPLEMENTARY DATA

Supplementary data are available at *JXB* online.

**Supplementary Fig. S1.** The dexamethasone (DEX)-inducible system to drive the expression of AtNEET, or its mutated dominant-negative copy H89C, in mature transgenic Arabidopsis plants.

**Supplementary Fig. S2.** Changes in steady state expression of different transcripts involved in reactive oxygen species (ROS) scavenging reported previously (Zandalinas et al., 2020b) in two different lines with constitutive expression of AtNEET or H89C.

**Supplementary Fig. S3.** Changes in protein expression associated with other functions of ferredoxins during the course of the experiment.

**Supplementary Fig. S4.** Changes in steady state expression of different transcript associated with iron-sulfur cluster assembly in the chloroplast and cytosol in three different homozygous H89C lines (H1, H7 and H9) following 4 doses of DEX application (Fig. 1A).

**Supplementary Table S1**. List of proteins altered following DEX treatment of Col and the inducible AtNEET and H89C lines.

**Supplementary Table S2.** Transcript-specific primers used for relative expression analysis by RT-qPCR.

## ACKNOWLEDGMENTS

Proteomic analyses were performed by The Charles W Gehrke Proteomics Center at the University of Missouri, Columbia, Missouri, USA (http://proteomics.missouri.edu).

## AUTHOR CONTRIBUTIONS

SIZ and RM conceptualized the project, SIZ conducted the experiments, SIZ and LS generated vectors and transgenic plants, SIZ, DGM-C, RN, and RM wrote the manuscript, RM and DGM-C obtained funding for the research.

## CONFLICT OF INTEREST

The authors declare no conflicts of interest.

## FUNDING

This work was supported by funding from the National Science Foundation (IOS-2110017, IOS-1353886, MCB-1936590, IOS-1932639), the Bond Life Sciences Early Concept Grant, and the Interdisciplinary Plant Group, and University of Missouri.

## DATA AVAILABILITY

The data supporting the findings of this study are available from the corresponding author, Ron Mittler, upon request. Proteomics data is deposited in the MASSIVE database (massive.ucsd.edu)with identifier PXD033795.

Reviewers can access via MSV000089456_reviewer with password SaraAtNEET51022.

---

**Supplementary Fig. S1.**
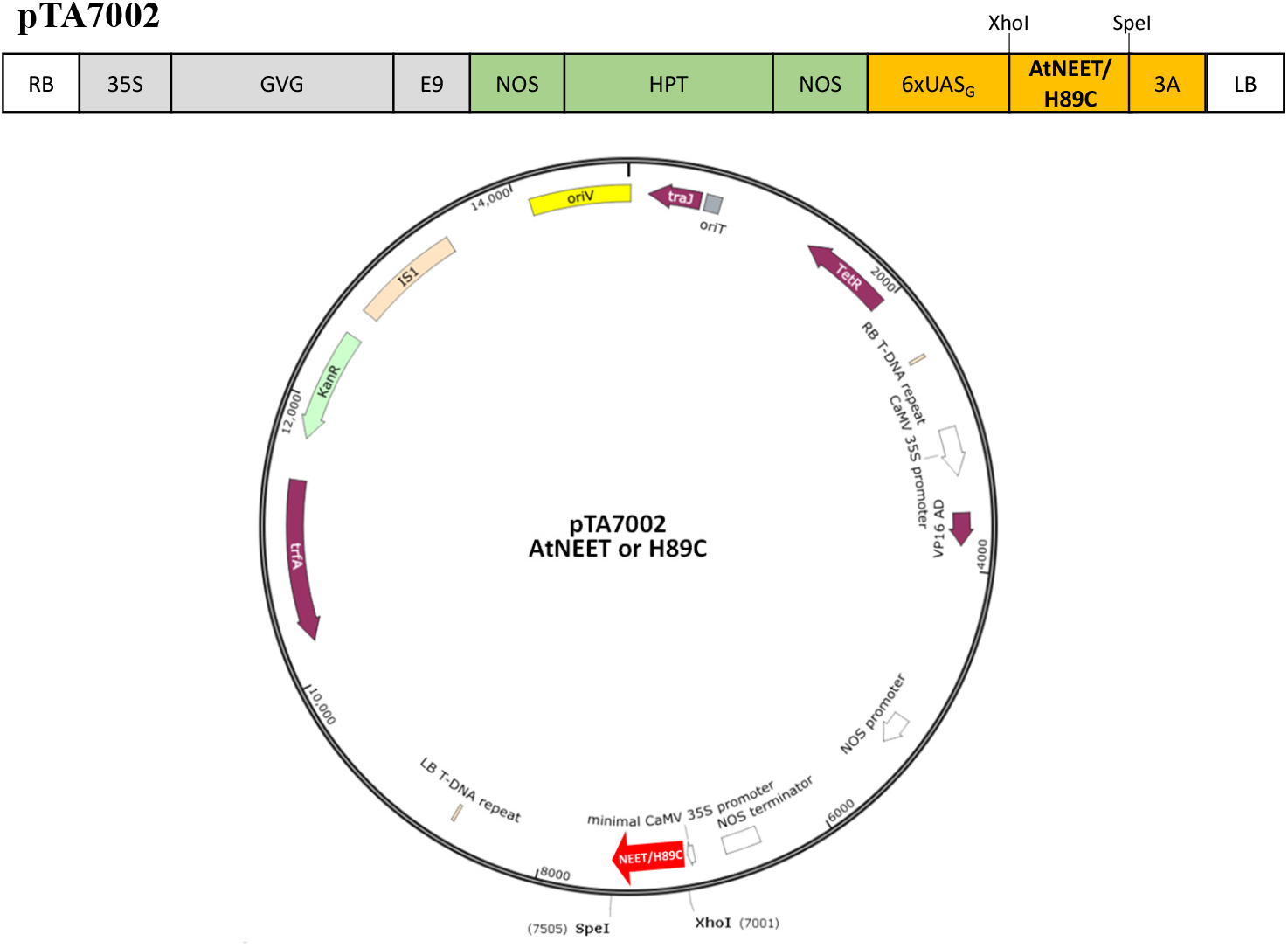
The dexamethasone (DEX)-inducible system to drive the expression of AtNEET, or its mutated dominant-negative copy H89C, in mature transgenic Arabidopsis plants. AtNEET or H89C were amplified and cloned into pTA7002 vector (Aoyama and Chua, 1997) using XhoI and SpeI sites. Abbreviations used: RB, right border; 35S, cauliflower mosaic virus 35S promoter; E9, the poly(A) addition sequence of the pea ribulose bisphosphate carboxylase small subunit rbcs-E9; GVG, chimeric transcription factor GVG composed by the heterologous DNA-binding of the yeast transcription factor <u>G</u>AL4, the transactivating domains from the herpes viral protein <u>V</u>P16, and the glucocorticoid receptor (<u>G</u>R); NOS, nopaline synthase promoter and terminator; HPT, hygromycin phosphotransferase; 6xUAS_G_, six copies of the GAL4 UAS; 3A, the poly(A) addition sequence of the pea rbcS-3A; LB, left border.

**Supplementary Fig. S2.**
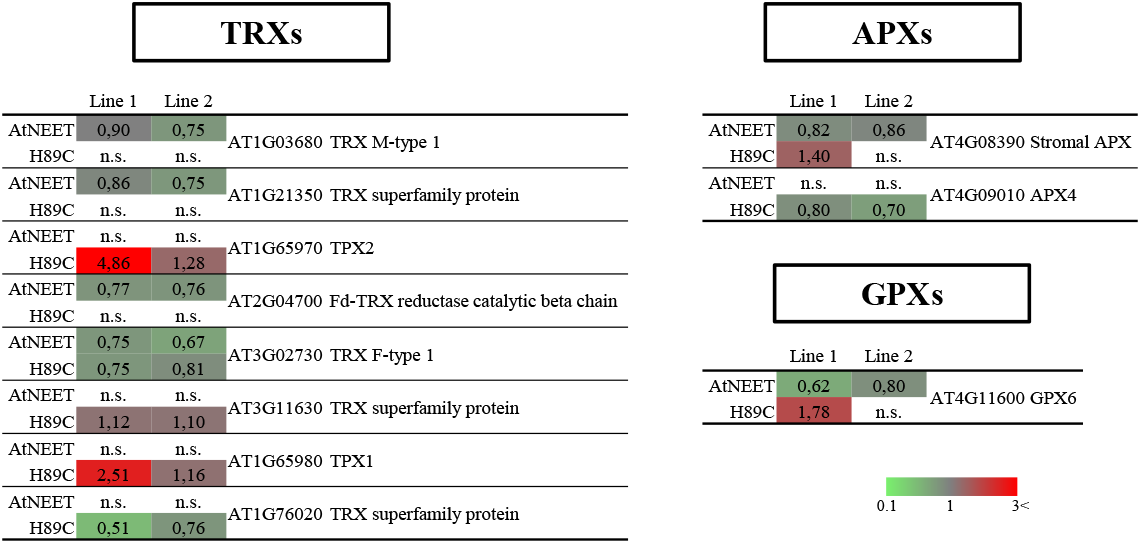
Changes in steady state expression of different transcripts involved in reactive oxygen species (ROS) scavenging reported previously (Zandalinas et al., 2020*b*) in two different lines with constitutive expression of AtNEET or H89C. Abbreviations used: APX, ascorbate peroxidase; GPX, glutathione peroxidase; n.s., not significant; TRX, thioredoxin.

**Supplementary Fig. S3.**
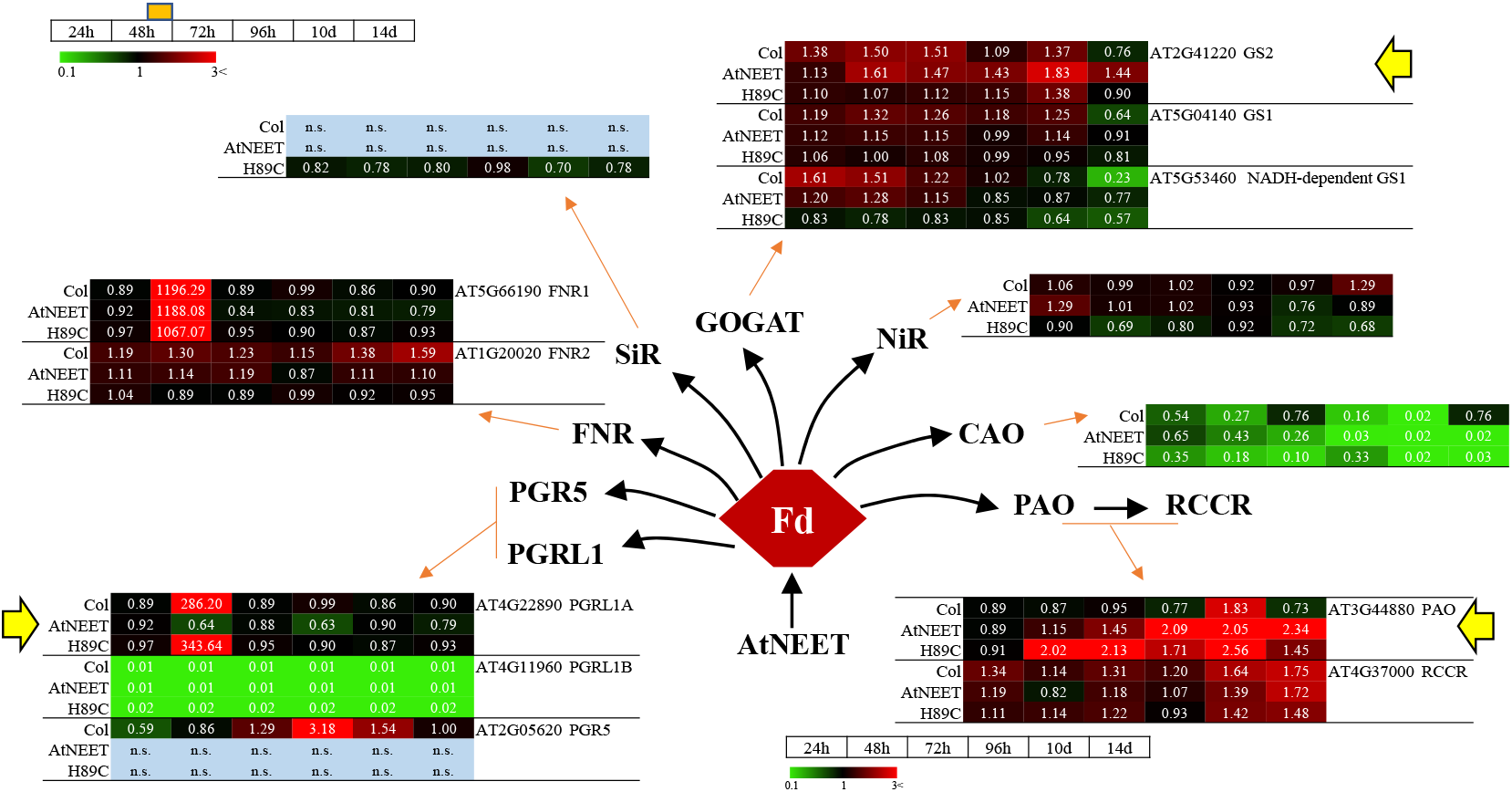
Changes in protein expression associated with other functions of ferredoxins during the course of the experiment. Pathway and heat maps for the expression of different proteins with a significant change in expression (in at least one time point, compared to time 0 h within each genotype) associated with additional ferredoxin functions in Arabidopsis at the different time points are shown. All experiments were repeated at least three times with similar results. Yellow arrows highlight proteins of interest. Abbreviations used: CAO, chlorophyll A oxygenase; Fd, ferredoxin; FNR, ferredoxin-NADP(+)-oxidoreductase; GOGAT, glutamine oxoglutarate aminotransferase; GS, glutamine synthase; NiR, nitrite reductase; PAO, pheophorbide A oxygenase, PGR5, proton gradient regulation 5; PGRL1A, proton gradient regulation 5-like A; PGRL1B, proton gradient regulation 5-like B; RCCR, red chlorophyll catabolite reductase; SiR, sulfite reductase.

**Supplementary Fig. S4.**
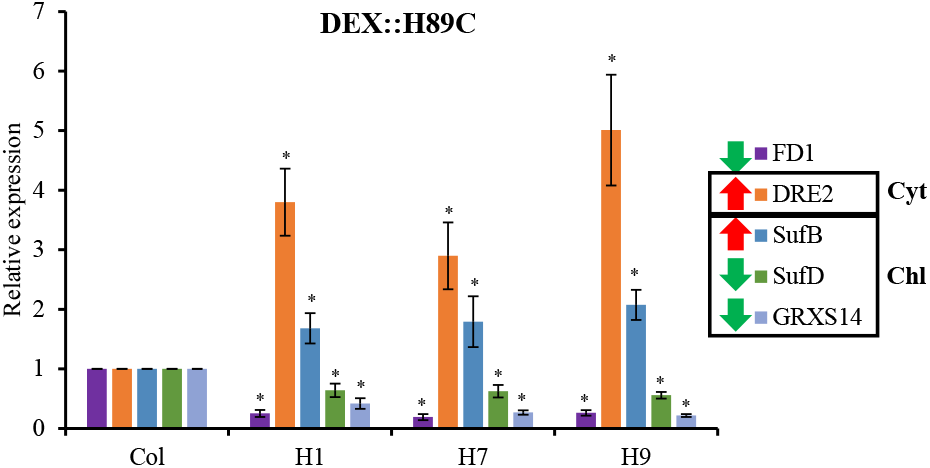
Changes in steady state expression of different transcript associated with iron-sulfur cluster assembly in the chloroplast and cytosol in three different homozygous H89C lines (H1, H7 and H9) following 4 doses of DEX application (Fig. 1A). All experiments were repeated at least three times with similar results. Asterisks denote statistical significance with respect to control (Col) at P < 0.05 (Student t-test, SD, N=5). Abbreviations used: Chl, chloroplast; Cyt, cytosol; DEX, dexamethasone; DRE2, Homolog of Yeast DRE2; FD1, ferredoxin1; GRXS14, Glutaredoxin S14; SufB, Sulfur B; SufD, Sulfur D.

## Notes

### Competing Interest Statement

The authors have declared no competing interest.

